# Allosteric activation of PARP2 self-PARylation by SUMO constrains DNA break-dependent catalytic Function

**DOI:** 10.1101/2025.08.29.673013

**Authors:** Arvind Parate Shivani, Eerappa Rajakumara

## Abstract

Poly(ADP-ribose)polymerase 2 (PARP2) is a key player in sensing DNA breaks and initiating DNA damage repair by catalysing the transfer of ADP-ribose units from NAD^+^ to target proteins, a process known as Poly(ADP-ribosyl)ation (PARylation). Post-translational modifications (PTMs) such as phosphorylation, ubiquitylation, SUMOylation, and PARylation are intricately linked to the DNA damage response (DDR) and repair. However, it is often overlooked that physical interactions between these enzymes and PTMs lead to DNA damage detection, DDR, and DNA repair. SUMOylation plays a vital role in DDR and DNA repair through covalent modification and non-covalent interactions. Here, we report new insight that Small ubiquitin like modifier (SUMO) binds with human PARP2 through non-covalent interactions, predominantly mediated by the N-terminal region (NTR) of PARP2. Surprisingly, SUMO stimulated PARP2 self-PARylation activity but hampered the DNA-dependent stimulation. Further competition binding studies suggest that SUMO binding promotes DNA release from PARP2. Altogether, our work uncovers a novel mechanism of SUMO-mediated allosteric regulation of PARP2 function, providing new insights into the possible interplay between SUMOylation and PARylation in DDR and DNA repair.

## Introduction

The cellular response to DNA damage involves a complex network of molecular mechanisms critical for maintaining genome integrity (1,2). DNA damage sites serve as hubs for the coordinated activation of signalling pathways and the recruitment of appropriate repair proteins (3). Among the key players in DNA damage response is Poly(ADP-ribose)polymerase 2 (PARP2), a nuclear enzyme primarily involved in sensing DNA strand breaks and initiating DNA repair (4–8). PARP2 is selectively activated by 5′ phosphorylated DNA breaks, suggesting its role in surveillance and resolution of pre-ligation repair intermediates to ensure proper DNA damage repair (4–8). PARP2 catalyses the transfer of ADP-ribose units from NAD^+^ to target proteins, a process known as PARylation, thereby facilitating the recruitment of DNA repair factors (4–8). PARP2 accounts for 5-15% of total cellular PARylation activity, along with PARP1 (75-95%), after DNA damage (9–14). Structurally, human PARP2 comprises: the unstructured N-terminal region (NTR), the Trp-Gly-Arg rich domain (WGR), the catalytic domain (CAT), which contains the catalytic ADP-ribosyl transferase (ART) domain (responsible for ADP-ribosylation), and a helical domain (HD) that serves as a regulatory domain for autoinhibitory function (4,6) (Fig. 1A).

**Figure 1:**
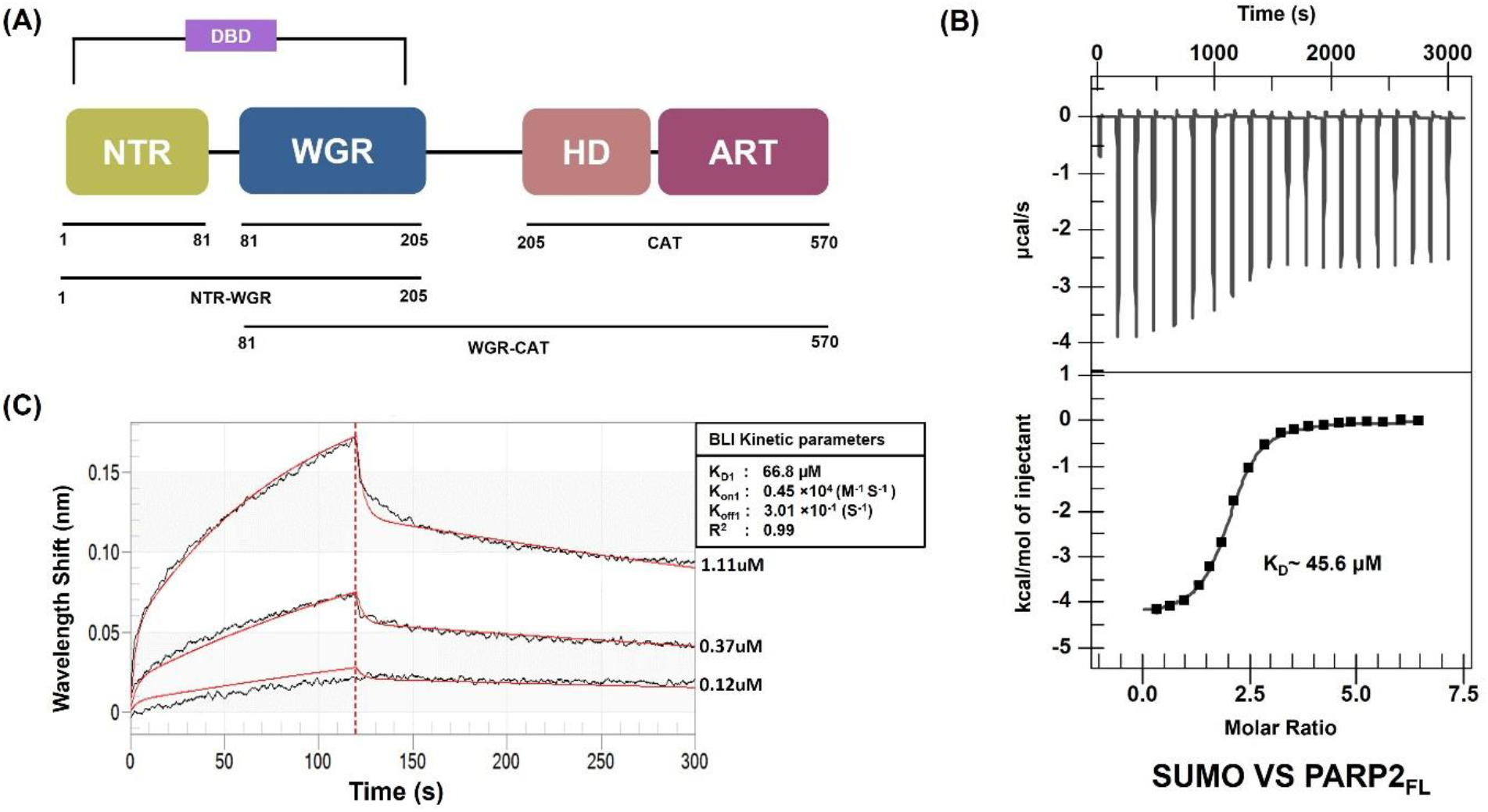
Measurement of binding affinity and kinetics of binding between PARP2 and SUMO. (A) Schematic representation of the domain architecture of human PARP2 (isoform 2) with black bars below the architecture representing the multi-domain constructs of PARP2 used in the study. NTR, WGR, CAT, HD, and ART stand for ‘N-terminal region’, ‘Tryptophan-Glycine-Arginine rich domain’, ‘catalytic domain’, ‘Helical domain’, and ADP-ribosyl transferase’, respectively. (B)ITC measurement of SUMO-PARP2_FL_ binding. Raw ITC data (upper panel) and normalized integration data (lower panel) for binding of SUMO to PARP2_FL_. (C)BLI sensogram of PARP2_FL_ binding to SUMO, where SUMO was biotinylated and immobilized on the biosensor. The data were fit globally to the 2:1 binding model. The processed experimental data curves are in black: model fit curves from the 2:1 global analysis are shown in red. R^2^ signifies the statistical confidence of the global fit of experimental data against model data.

Post-translational modifications (PTMs) of chromatin and chromatin-associated proteins provide a rapid and reversible means to label DNA lesions and their surrounding chromatin landscape (15–17). PTMs are key in controlling the dynamic molecular architecture of DNA repair and signalling proteins (15,16). These modifications, such as Phosphorylation, Ubiquitylation, SUMOylation, and PARylation are intricately linked to the DNA damage response (DDR) (18–20). However, it is often overlooked that communication between these enzymes and PTMs, through physical interactions, occurs during the DNA damage response (DDR), a cellular network that detects, signals, and repairs DNA damage to maintain genomic integrity.

SUMOylation is a critical regulatory mechanism that influences various DDR pathways by modulating protein-protein interactions, protein stability, and localization (2,20). SUMO conjugation observed in response to DNA breakage promotes the accumulation of ubiquitin chains on damaged chromatin and is required for the efficient recruitment of ubiquitin-dependent genome caretakers (20,21). SUMOylation not only contributes to the recruitment of proteins to DSBs but also their coordinated removal, and interestingly, is required for both non-homologous end joining (NHEJ) and homologous recombination (HR), in which PARP2 also plays a significant role (2,14,22–24). SUMO interacts non-covalently with target proteins via SUMO-interacting motifs (SIMs) or other recognition elements, adding another layer to the regulatory network (2,25).

Previous studies have demonstrated functional intersections between SUMO and PARP1 in the DNA damage response (2,19,26,27). Notably, PARP1 is SUMOylated by the E3 ligase PIAS4 and regulated through ubiquitination and acetylation (27). However, the possible direct, non-covalent interaction between SUMO and PARP2 remains poorly understood, and the contribution of specific PARP2 domains to SUMO binding has not been explored. Understanding whether or how SUMO recognizes PARP2 and modulates PARP2 activity could uncover novel regulatory mechanisms in DNA repair and chromatin dynamics.

In this study, we employed biophysical and biochemical approaches to investigate the interaction between PARP2 and SUMO. Our data reveal that human PARP2 can recognize SUMO through a direct, non-covalent interaction. Domain mapping studies indicate that the N-terminal region (NTR) of PARP2 significantly contributes to this interaction, suggesting a potential SUMO-binding interface within this region. Enzymatic assays further demonstrate that SUMO binding enhances the catalytic activity of PARP2. Interestingly, SUMO appears to function as an allosteric regulator, capable of modulating PARP2 activity in both DNA-dependent and independent contexts. Further, SUMO displaces DNA break from the PARP2-DNA break complex, which hampers the DNA breaks-dependent catalytic activity of PARP2. These findings suggest that SUMO serves as a docking site for PARP2 and plays a nuanced role in fine-tuning the catalytic output of PARP2, potentially influencing the temporal and spatial dynamics of PARP2 mediated signalling during the DNA damage response (DDR).

## Results and discussion

### Biophysical characterization of SUMO binding to PARP2

SUMO modulates the function of target proteins through covalent attachment and non-covalent interactions through SUMO interacting motif on the corresponding target protein (2,28). Studies have shown that the DNA repair proteins, such as XRCC4, SLX4, SETDB1, and Srs2 interact with SUMO through its SUMO-interacting motif (SIM), highlighting a broader regulatory role for non-covalent SUMO-protein associations (23,29–31). Given that many proteins involved in the SUMOylation pathway also engage in non-covalent interactions with SUMO, we hypothesized that PARP2, an essential DDR mediator, may also bind to SUMO.

To test this hypothesis, we employed isothermal titration calorimetry (ITC) binding studies to quantify the binding affinity (K_D_) of the SUMO–PARP2 interaction using the recombinant SUMO and PARP2_FL_ (Supplementary Fig. 1). Our ITC data revealed that SUMO binds to PARP2_FL_ with a K_D_ of approximately 45.6µM, indicating a moderate binding affinity (Fig. 1B).

We conducted biolayer interferometry (BLI) using biotinylated SUMO and purified recombinant PARP2_FL_ (Supplementary Fig. 1) to validate and determine kinetic parameters of this interaction. The BLI analysis confirmed a similar equilibrium dissociation constant (K_D_∼66.8 µM) (Fig. 1C). Kinetic analysis further yielded an association rate constant (k_on_) of ∼0.45 × 10^4^ M^−1^S^−1^ and a dissociation rate constant (k_off_) of ∼3.01 × 10^−1^ S^−1^, validating the formation of a moderately stable complex between SUMO and PARP2_FL_. The micromolar K_D_ value, coupled with the relatively slow k_off_ rate (suggesting high residence time), supports the formation of a specific and stable PARP2_FL_-SUMO complex.

### SUMO - PARP2 Association Is Driven by the Intrinsically Disordered N-Terminus of PARP2

Following these findings, we further examined which domains of PARP2_FL_ are responsible for SUMO recognition to elucidate the molecular basis of this interaction. The N-terminal region (NTR) of the PARP2 is intrinsically disordered and highly enriched in basic amino acids, particularly lysine and arginine residues, which confer a strong positive charge and facilitate electrostatic interactions with negatively charged protein surfaces (5,14,32).

The isoelectric point (pI) of the NTR is ∼11.0, which supports the above observation. Previous studies have shown that the N-terminal region plays an essential role in the binding of negatively charged regulators like DNA (12,33). SUMO, enriched in acidic patches and possessing a pI of ∼5.9, can interact with lysine or other positively charged residues within the intrinsically disordered regions of proteins, making them favourable partners for electrostatic engagement (28). Therefore, assuming that these two interact through ionic interactions, we first analysed the binding between the recombinant SUMO and the N-terminal region of PARP2 (NTR_PARP2_) (Supplementary Fig. 1). The ITC binding studies using the recombinant SUMO and NTR_PARP2_ revealed that NTR_PARP2_ exhibited measurable binding affinity (K_D_∼ 201µM) (Fig. 2A).

**Figure 2:**
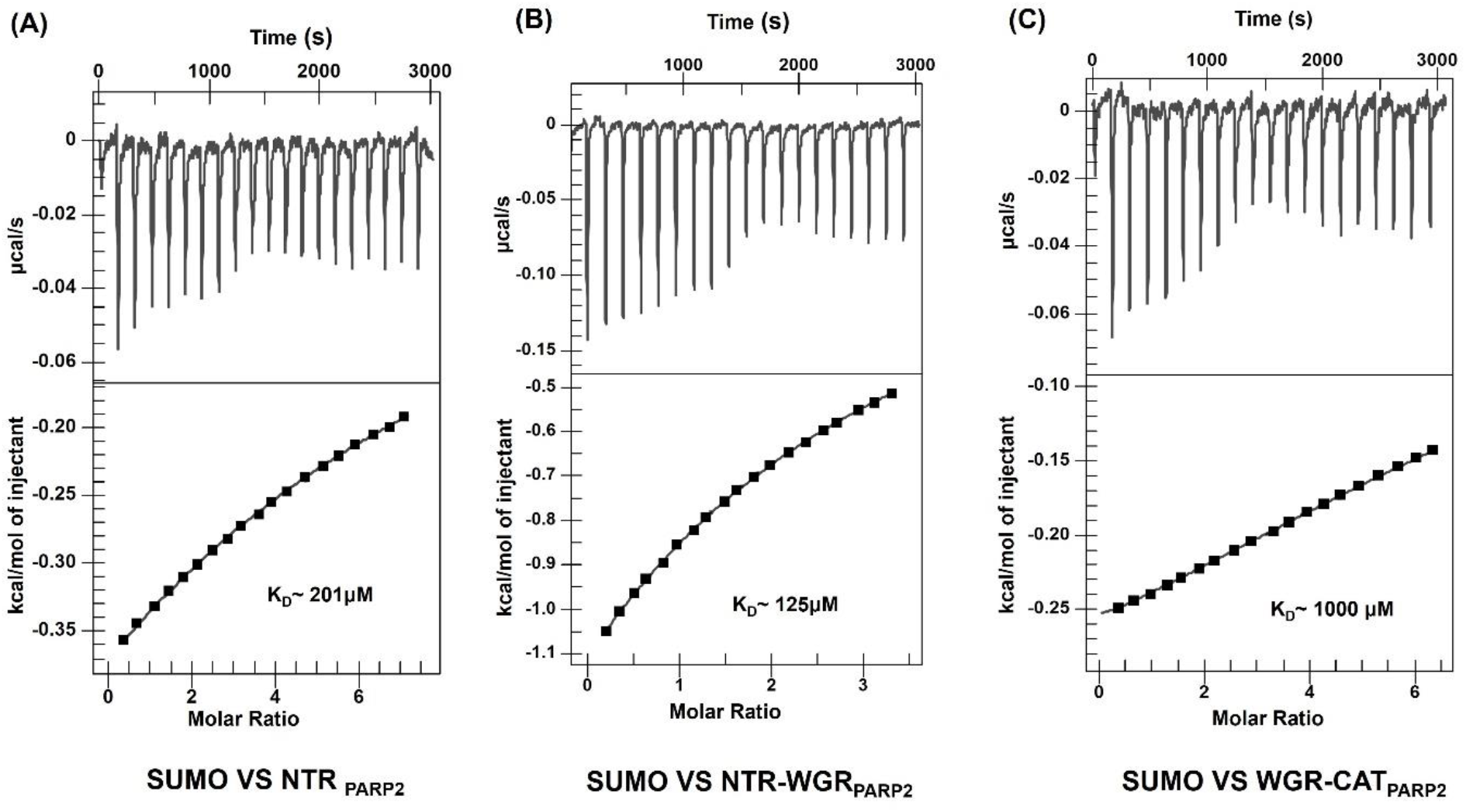
ITC measurement of the binding of SUMO with domains of PARP2. Raw ITC data (upper panel) and normalized integration data (lower panel) for binding of SUMO to (A) NTR_PARP2_, (B) NTR-WGR_PARP2_ and (C) WGR-CAT_PARP2_.

A weak binding affinity between the NTR_PARP2_ and SUMO suggested that the WGR and CAT domains may mainly contribute to the binding affinity between the SUMO and PARP2. Surprisingly, recombinant WGR_PARP2_ and CAT_PARP2_ domains (Supplementary Fig. 1) alone did not exhibit measurable binding towards SUMO (Supplementary Fig. 2C and D). This observation underscores the dominant role of the NTR in SUMO recognition.

### SUMO Recognition mediated by different domain combinations of PARP2

The N-terminal region (NTR) and WGR domain of PARP2 are known to sense PARP2 activators and induce conformational changes in the catalytic domain, thereby activating PARP2-mediated PARylation (12,34). The above binding studies suggested that combining NTR with other domains might contribute to the observed binding affinity between the SUMO and PARP2. Next, we assessed domain combinations to understand cooperative contributions. The recombinant NTR-WGR_PARP2_ (Supplementary Fig.1) construct displayed a slightly higher K_D_ (∼125 µM) than NTR_PARP2_ alone, highlighting the dominant role of the N-terminal domain and a significantly weaker contribution of WGR in SUMO binding (Fig. 2B). The recombinant WGR-CAT_PARP2_ (Supplementary Fig. 1) also bound SUMO with a considerably weaker affinity (K_D_ ∼1000 µM), approximately 5-fold and 22-fold lower compared to NTR_PARP2_ and PARP2_FL_, respectively (Fig. 2C). From the results of the above binding studies, we concluded that entire PARP2 contributed for NTR-driven binding to SUMO.

### SUMO-Mediated Regulation of PARP2 Catalytic Function

Following the confirmation of PARP2_FL_–SUMO interaction (Fig. 1B and C), we investigated whether SUMO influences the catalytic activity of PARP2 through an SDS-PAGE-based in vitro auto-modification assay (35–37) using PARP2_FL_ complexed with increasing concentrations of SUMO. The loss of the PARP2_FL_ band intensity and the appearance of a higher molecular weight smear indicated auto-PARylation activity, suggesting that SUMO enhances the catalytic activity of PARP2_FL_ (Fig. 3A; lanes 5–9). As a positive control, we employed 5′ phosphorylated single-strand break (5′P-SSB) (Table S1), a well-characterized allosteric activator of PARP2 (4–8), which resulted in robust catalytic activation (Fig. 3A; lane 4). Similarly, 5′ phosphorylated double-strand break (5′P-DSB) (Table S1) was also used as a control and exhibited maximal PARP2_FL_ activation (Fig. 3B; lane 4). An increase in SUMO concentration (Fig. 3B; lanes 5–9), correlating with an intensified smear above the PARP2_FL_ band, further confirmed SUMO-mediated stimulation of PARP2_FL_ auto-modification activity. This property of the PARP2 suggests that SUMO-dependent stimulation of enzymatic activity may be required for the PARylation of proteins involved in DNA repair and other nuclear processes.

**Figure 3:**
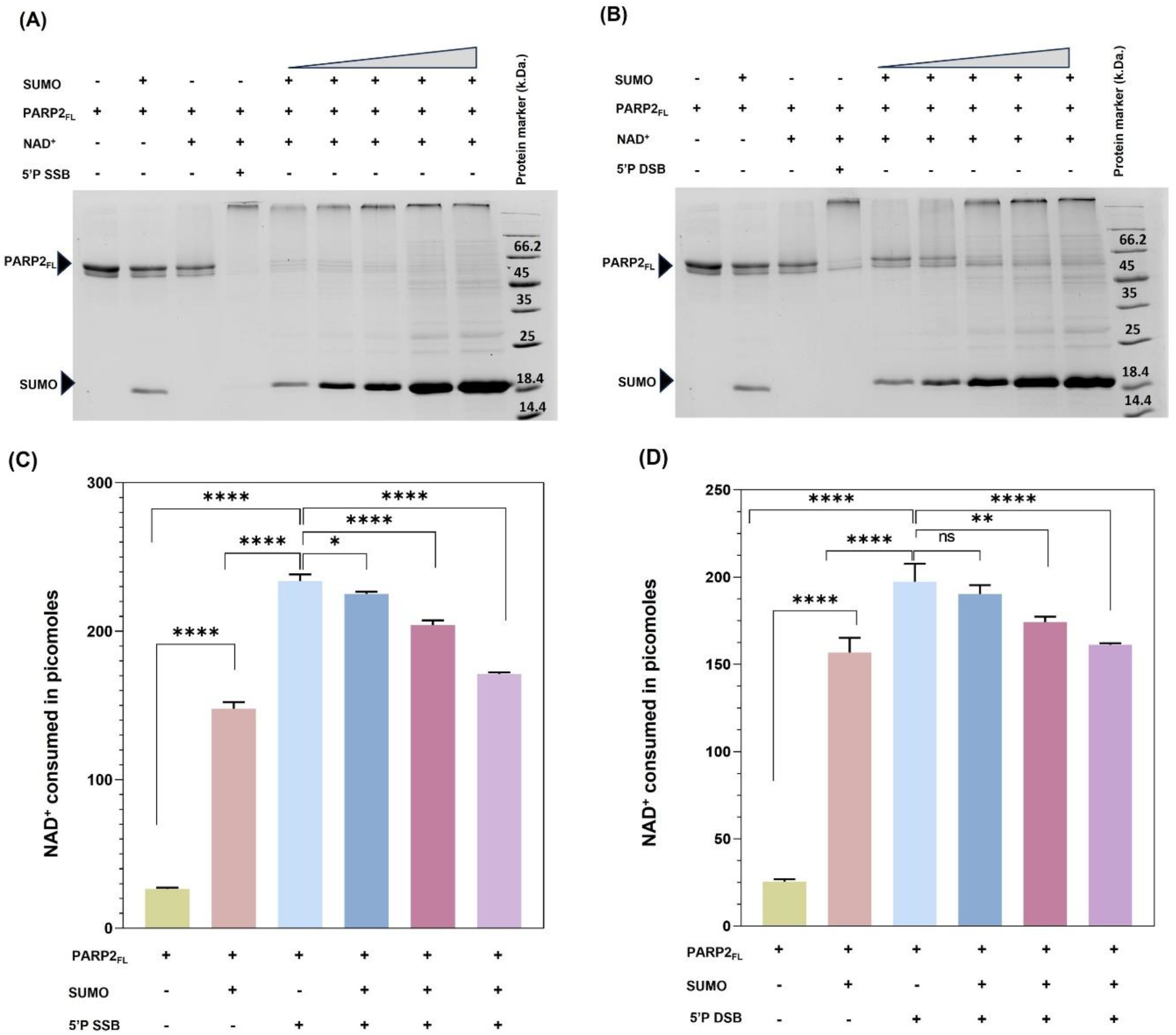
PARP2 Auto-modification Assay. (A) SDS-PAGE PARylation assay of PARP2_FL_ in the presence of 5′P-SSB DNA and increasing concentration of SUMO. (B) SDS-PAGE PARylation assay of PARP2_FL_ in the presence of 5′P-DSB DNA and increasing concentrations of SUMO. (C) Quantification of PARylation activity of PARP2_FL_ in the presence of 5′P-SSB DNA, SUMO, and a combination of 5′P-SSB DNA and SUMO. (D) Quantification of PARylation activity of PARP2_FL_ in the presence of 5′P DSB DNA, SUMO, and a combination of 5′P-DSB DNA and SUMO. PARylation activity of PARP2 was determined using a Acetophenone-KOH based chemical method. The p-values were calculated using One Way Anova test where *, **, *** and **** indicates P-values < 0.05, < 0.005, < 0.001 and < 0.0001 respectively. Error bars indicate the standard deviation of the n = 3 independent experiments.

To assess the extent of PARylation, we performed the acetophenone-KOH based chemical assay, which measures residual NAD^+^ following the PARylation reaction (33,37–42). In this assay, acetophenone and KOH react with unconsumed NAD^+^ in the presence of formic acid to generate a fluorescent product, the intensity of which is directly proportional to the concentration of unused NAD^+^. As reported previously, a standard curve was used to determine NAD^+^ concentrations in the reactions (37). Reactions were conducted with 10μM NAD^+^ at room temperature for 5 minutes. The in-vitro PARylation assay revealed that PARP2_FL_ exhibited ∼5.5-fold and ∼8.4-fold higher catalytic activity in the presence of SUMO and 5′P-SSB, respectively, compared to basal activity (Fig. 3C). A similar assay using 5′P-DSB as the DNA activator showed ∼6.2-fold enhancement of PARP2_FL_ activity in the presence of SUMO and ∼7.7 -fold stimulation in the presence of 5′P-DSB alone. (Fig. 3D)

### Effect of SUMO on DNA-dependent stimulation of PARP2 catalytic activity

Several allosteric ligands combinatorially modulate the receptor’s activity to fine-tune the functional outcome (35,36,43,44). Given that both DNA breaks (4–8) and SUMO (Fig. 1B and C; Fig. 3) bind and stimulate the PARP2 catalytic activity, here we set out to investigate whether SUMO binding to PARP2 regulates the DNA-break-dependent catalytic activity of PARP2 using the acetophenone KOH-based chemical assay, (33,37–42). We quantified the extent of DNA breaks dependent stimulation of catalytic activity in the presence of an increasing concentration of SUMO. We observed the DNA-dependent catalytic activity of PARP2_FL_ marginally decreased with 1:1 molar ratio of DNA-break and SUMO (Fig. 3C and D). However, as the concentration of SUMO increased (1:3; 1:5 molar ratio), PARP2_FL_ activity progressively declined (Fig. 3C and D). This inverse relationship suggests SUMO hampers DNA-breaks-dependent catalytic activity of PARP2_FL_. These findings imply that SUMO can modulate PARP2 activity by engaging with the PARP2’s overlapping and/or distinct regions. Notably, the extent and nature of SUMO-mediated stimulation differ from the classic DNA-dependent activation, indicating that SUMO acts as a fine-tuning regulator of PARP2 in both DNA-dependent and -independent contexts.

### SUMO-Mediated Displacement of DNA from PARP2

NTR-WGR of PARP2 binds to DNA breaks (12,34), and the current studies suggest that NTR-WGR is a major contributor to SUMO recognition by PARP2 (Fig. 2B). Further, SUMO hampers the DNA-breaks-dependent catalytic activity of PARP2. To investigate whether SUMO competes with DNA breaks binding to PARP2, we performed BLI-based competition binding studies. A 5′ phosphorylated biotinylated DSB (Table S1) DNA was immobilized on streptavidin-coated sensors, which were then dipped into solutions containing a PARP2_FL_–SUMO binary complex with an increase in SUMO concentrations from 1:0 to 1:16 molar ratio. If SUMO and DNA simultaneously bind to PARP2_FL_, one would expect an increased wavelength shift with the PARP2_FL_–SUMO complex compared to PARP2_FL_ binding to DNA alone. However, contrary to this expectation, the highest wavelength shift was observed when PARP2_FL_ alone bound to DNA (Fig. 4A). The addition of SUMO progressively reduced the binding signal. At a SUMO: PARP2_FL_ ratio of 16:1, no significant wavelength shift was detected, indicating that increasing SUMO levels displace DNA from the PARP2_FL_ complex, hindering its DNA-binding ability, especially in the context of 5′P-DSB DNA. Comparing the BLI signals at report points at the end of the association phase with the absence of competitor (SUMO) allowed us to quantify the competition for every SUMO concentration, and plotting the respective nm values (shifts) against SUMO concentrations gave an IC_50_ value of 0.82μM (Fig. 4B).

**Figure 4:**
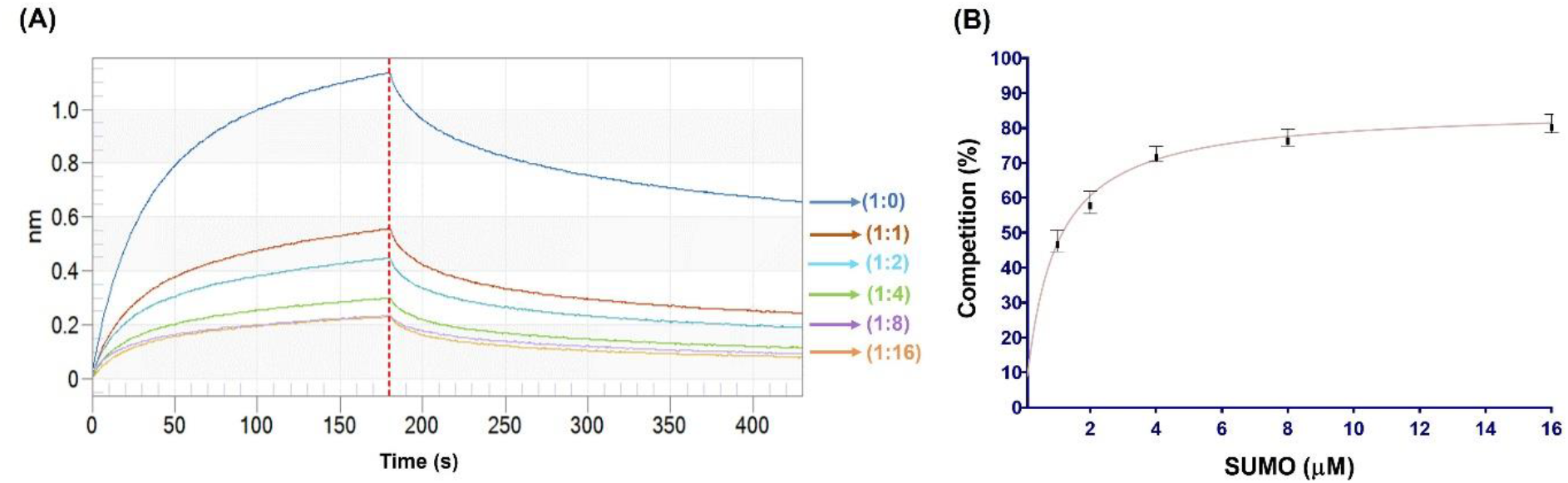
BLI Competition binding studies: (A) Representative sensogram showing a progressive decrease in wavelength shift (binding signal) with increasing SUMO concentration (1µM to 16µM), indicating reduced association of the PARP2–SUMO complex with immobilized 5′P-DSB DNA (B) Corresponding dose response curve from BLI experiment measuring concentration dependent competition of SUMO and 5′P-DSB to bind with PARP2. The IC_50_ value was determined by nonlinear regression using a 3-parameter logistic fit with variable slope. Data represent the mean and standard deviation of triplicate measurements.

## Conclusion

In conclusion, this study uncovers a previously uncharacterized, direct, non-covalent interaction between human PARP2 and SUMO, providing new insights into the allosteric regulation of PARP2. Using biophysical and biochemical assays, we demonstrate that the intrinsically disordered, positively charged N-terminal region (NTR) of PARP2 serves as a key interface for SUMO binding. While this interaction alone exhibits moderate affinity, the cooperative contribution of additional domains within full-length PARP2 enhances binding stability. Functionally, SUMO binding stimulates PARP2’s catalytic PARylation activity in both DNA-dependent and - independent contexts. However, at higher concentrations, SUMO competes with DNA for PARP2 binding, thereby attenuating its DNA-break-stimulated activity. These findings highlight SUMO as a dual-function regulator of PARP2, acting both as an activator and a competitive modulator of DNA break-dependent allosteric activity of PARP2. This interplay between SUMO and PARP2 might add a critical layer of regulation to DDR signalling, influencing the spatial and temporal dynamics of PARP2 activity. Further structural studies of PARP2-SUMO complex would reveal the structural basis of allosteric activation of PARP2 catalytic activity, similarities and differences with DNA break-dependent allosteric activation, and the mechanistic insights into competition with DNA breaks binding to PARP2, and constraining DNA break-dependent allosteric stimulation of PARP2 activity.

## Materials and Methods

### Cloning, Expression and Purification of hPARP2_FL_, WGR-CAT_PARP2,_ WGR_PARP2_, NTR_PARP2_, NTR-WGR_PARP2_

Full-length human PARP2 isoform 2 (amino acids 1–570), hereafter referred to as PARP2_FL_ (NCBI Accession: NP_001036083), was kindly provided by Dr. John Pascal in the pET28a (+) expression vector. The following constructs were generated, including the N-terminal region (NTR; aa 1–80), WGR domain (aa 81– 205), WGR-CAT (aa 81–570), and the NTR-WGR construct (aa 1–205). These fragments were cloned into the pRSFDuet-1 vector using *BamHI* and *XhoI* restriction sites. A description of the domain length of individuals and the combination of the domains is given in (Fig 1A). All constructs were transformed into *E. coli* Rosetta2 (DE3) cells for expression. An overnight primary culture was prepared by picking colony from the transformed plate in LB medium containing 10mM benzamide, 50µg/mL kanamycin, and 35µg/mL chloramphenicol, and grown at 37 °C. This primary culture was used to inoculate a secondary culture, which was grown until the OD_600_reached 0.8 approximately. Protein expression was induced with 0.2mM IPTG, and the culture was supplemented with 10mM benzamide, followed by incubation at 16 °C for 16–18 hours. Cells were harvested by centrifugation at 8000 × *g* for 10 minutes at 25 °C. Cell pellets were resuspended in lysis buffer containing 500mM NaCl, 50mM HEPES (pH 7.5), 20 mM imidazole, 3mM β-mercaptoethanol, and 1mM phenylmethylsulfonyl fluoride (PMSF). Cell lysis was carried out by sonication, and the lysate was clarified by centrifugation at 10,500 rpm for 90 minutes at 4 °C. Protein purification was conducted in three steps: (1) Ni-NTA affinity chromatography using a 5 mL HisTrap column (GE Healthcare), (2) Heparin affinity chromatography using a HiTrap Heparin HP column with a NaCl gradient ranging from 250 to 1000mM, and (3) Gel filtration chromatography using a Superdex 200pg 16/600 column (GE Healthcare), pre-equilibrated with buffer containing 150mM NaCl, 10mM HEPES (pH 7.5), 10% glycerol, and 2mM β-mercaptoethanol as reported previously (35,37,45). The final protein fractions were concentrated, flash-frozen in liquid nitrogen, and stored at – 80 °C. The same purification strategy was applied to all PARP2_FL_ and its domains. The SDS-PAGE gel images of all purified proteins are provided in (Supplementary Fig.1A-F).

### Expression and Purification of SUMO and ULP1

The Champion™ pET SUMO Expression System was used for production of SUMO protein. The Champion™ pET SUMO vector and in house cloned ULP1 construct in the pRSF-DUET1 expression vector were transformed into *E. coli* BL21(DE3). An overnight primary culture was prepared by picking a colony from the transformed plate in LB medium containing 50µg/mL kanamycin and grown at 37 °C. This primary culture was used to inoculate a secondary culture, which was grown until the OD_600_reached 0.8 approximately. Protein expression was induced with 0.2mM IPTG, and followed by incubation at 16 °C for 16–18 hours. Cells were harvested by centrifugation at 8000 × *g* for 10 minutes at 25 °C. Cell pellets were resuspended in lysis buffer containing 500 mM NaCl, 50mM HEPES (pH 7.5), 20mM imidazole, 3mM β-mercaptoethanol, and 1 mM phenylmethylsulfonyl fluoride (PMSF). Cell lysis was carried out by sonication, and the lysate was clarified by centrifugation at 10,500 rpm for 90 minutes at 4 °C. ULP1 was purified, employing Ni-NTA affinity chromatography with a 5 mL HisTrap column (GE Healthcare). Further SUMO protein was purified by Ni-NTA affinity chromatography as described earlier (35,37,45). Ni-NTA eluates were incubated overnight with an in house ULP1 and then protein was subsequently purified using the Mono Q anion exchange chromatography column, followed by the gel filtration chromatography to get pure SUMO protein. The SDS-PAGE gel images of all purified proteins are provided (Supplementary Fig. 1G and H).

### Isothermal titration calorimetry binding studies

Isothermal titration calorimetry binding studies were conducted at 25 °C in LV Affinity ITC (TA Instruments, New Castle, DE, USA). The reference cell was filled with deionized water. For all binding studies, unless stated otherwise, PARP2_FL_ (70μM) and PARP2 domains (70μM) were taken in the sample cell (350μL) and SUMO (1– 1.5 mM) was taken in the syringe. Two consecutive injections of 2.5μL for a total 20 injections at an interval of 120s were performed while stirring the cell content at 125 rpm. As a control, the SUMO (1mM) was titrated against buffer and as well as buffer was titrated against PARP2 constructs (70μM) (Supplementary Fig. 2A and B). For sample preparation, titrants and titrands were diluted in gel filtration buffer (150mM NaCl, 10mM HEPES, 2% Glycerol, and 2mM β-mercaptoethanol). All the binding studies were repeated three times. Analysis and processing of the binding isotherm were performed using NANOANALYZE software provided by TA instruments, and the ITC data was deconvoluted in “independent” model of curve fitting using a nonlinear least-squares algorithm.

### Generation of biotinylated SUMO

Biotin labelling of SUMO was done by biotinylating reagent i.e. EZ-Link NHS-LC-LC-Biotin (Thermo Scientific) (44). First 10mM of Stock solution of NHS -LC-LC linker was prepared in 350μl of DMSO, then sub stock of 1mM biotinylating reagent was prepared in 1000μl of nuclease free water. Minimum protein concentration required for biotinylating is 1mg/ml. SUMO (2.6mg/ml) was incubated with 11.6μl of 1mM biotin reagent in 1:1 Molar coupling ratio in 50μl gel filtration buffer (50mM HEPES pH-8), 250mM NaCl, 2mM β-mercaptoethanol and 10% glycerol) for 2 hours at 4°C. The reaction mixture was desalted on a Zeba Spin Desalting Columns (Thermo scientific, 7k MWCO). Column preparation was done by removing the storage solution by centrifuging it at 1500g for 1 min. The reaction mixture was loaded on the column and incubated for 2 min so that resin absorbs the sample. The reaction mixture was centrifuged at 1500g for 2 min at 25°C and collected in microcentrifuge tube. Concentration was calculated by taking absorbance at 280 nm using molar extinction of SUMO (0.133M^-1^CM^-1^).

### Biolayer interferometry Binding Studies

Biolayer interferometry-based binding kinetics studies were carried out on the OctetK2 instrument (Pall Fortebio). Biotinylated ligand i.e. SUMO (15μg/ml) was immobilized onto High Precision Streptavidin (SAX) Dip and Read™ biosensors (Pall Fortebio, Fremont, CA, USA). For association kinetics measurement, the probes were dipped into wells containing varying concentrations of PARP2_FL_ (3-fold serial dilution;10μM-41nM) in buffer (150mM NaCl, 25mM Tris-HCl, 5% glycerol, 0.1% BSA (w/v), 0.01% Tween-20 (v/v) and 3mM β-mercaptoethanol for 120-150s followed by dipping into wells containing buffer for 180s to measure dissociation kinetics. For reference, one of the probes was dipped into buffer during the association and dissociation phases. The measurement was run at 25°C.The data were analysed and fit to 2:1 model using ForteBio Data Analysis 12 (Pall Fortebio) (Fig. 1B).

### Nucleic acids generation and annealing

HPLC-purified lyophilized oligonucleotides and complementary strands (listed in Supporting Table S1) were purchased from the oligo synthesis facility of W.M. Keck Foundation, Yale School of Medicine, USA. The oligos were dissolved in a buffer containing 10mM Tris-Cl (pH 7.5), 50mM NaCl, and 3mM MgCl_2_. 16 mer 5′P-DSB DNA were prepared by mixing the complementary strands in an equimolar ratio. A dumbbell-forming 45-mer DNA oligonucleotide was used to make 5′P-SSB DNA. 2mM of 5′P-SSB and 5′P-DSB DNAs were annealed as reported previously (35,46).

### SDS-PAGE based automodification Assay of PARP2

The automodification reactions of PARP2 were performed as reported earlier (35,36,38,39) with some modifications. Briefly, 1μM of PARP2_FL_ was incubated with equal molar ratio of 5′P-DSB and 5′P-SSB DNA and increasing concentration of SUMO (1μM to 10μM) in auto modification buffer (10mM HEPES, pH 8, 25mM KCl, 5mM CaCl_2_, 5mM MgCl_2_, and 2mM β-ME) accounting for 12μl of total reaction volume for 10 min to allow for possible protein−protein/protein−dna interactions to occur. Following the incubation, 2mM NAD^+^ was added in the reaction. The automodification reaction was kept at room temperature for 2 hours. After the indicated time point, the reactions were stopped by adding 4X SDS-loading buffer. The reaction solution was resolved on 15% SDS-PAGE. The gel was stained with Coomassie Blue dye. The increase in the molecular weight due to PARP2_FL_ auto-PARylation was identified in the gel as a smearing pattern extending upwards from a thin band corresponding to the left-over unmodified PARP2. Gel Doc (iBrightCL1000) was used to capture gel images (Fig. 3A and B).

### Quantification of PARP2 Activity

Activity assays were performed with minor modifications based on a previously published protocol (33,37–42). Reactions were carried out in 96-well microplates (Greiner Bio-One, PP, F-bottom) at room temperature (RT). Total reaction volume was 60µL which contained 1µM PARP2_FL_ protein, 1µM SUMO, and/or 5′P-DSB/SSB

DNA, along with 10µM NAD^+^. The reaction mixtures were incubated at RT for 5 minutes. Following incubation, 20µL of 20% acetophenone (prepared in absolute ethanol) and 20µL of 2M KOH were added to each well and mixed thoroughly. The plates were incubated at 4 °C for 10 minutes. Subsequently, 90µL of 88% formic acid was added, and the mixtures were heated at 110 °C for 5 minutes to develop fluorescence. The plates were then cooled to RT for 30 minutes. Fluorescence was measured using an EnSpire™ Multimode Plate Reader (PerkinElmer, Inc.), with monochromators set to an excitation wavelength of 360 nm and emission at 445 nm. The assay quantifies the remaining NAD^+^ in solution. Net fluorescence was calculated using Equation (1a), and the percentage of NAD^+^ consumption by PARP2_FL_ was determined using Equation (1b).

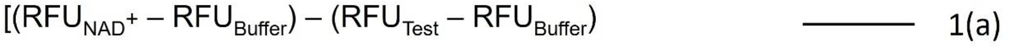

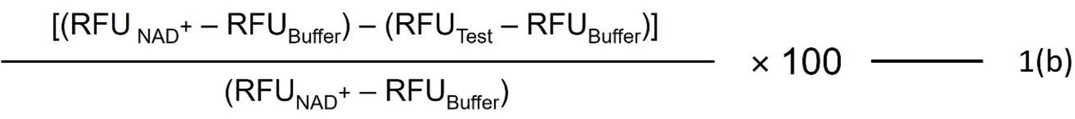

In equations (1a) and (1b), RFU_NAD_^+^ stands for Relative Fluorescence Unit with NAD^+^ only, RFU_Test_ stands for Relative Fluorescence Unit with tested sample, and RFU_Buffer_ stands for Relative Fluorescence Unit with buffer only. The statistical analysis was performed via ordinary One-way ANOVA using Graph Pad Prism software.

### Competition binding studies

To investigate the effect of SUMO on the PARP2-DNA interaction, we employed BLI binding studies using biotinylated 5′P-DSB DNA (1μg/mL) immobilized on the SAX biosensor (35,47). To get the association data, the biosensor was dipped in a well containing PARP2_FL_ (1μM) and then subsequently dipped in wells containing PARP2_FL_-SUMO complex with increasing concentration of SUMO from 1:0 to 1:16 molar ratios. The association was allowed to happen for 200 s. The sensor bound with DNA was regenerated between every association step by dipping the sensor in the regeneration buffer containing 5M NaCl, followed by a 180s dissociation step in a tube filled with assay buffer. All the samples were prepared in the same buffer used for BLI binding kinetics measurement. The experiment was run at 25°C. The data were analysed using ForteBio Data Analysis 12 (Pall Fortebio) (Fig. 4A). The binding signals at report points after 200s of association (B_M_) were used to calculate the degree of competition (C) for every ligand concentration compared with positive controls (B_P_) by using the following equation (47).

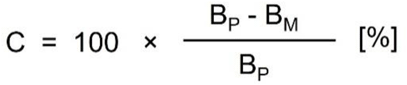

The calculated competition values were plotted against SUMO concentrations (μM), and the data were fit using non-linear regression to fit a sigmoidal dose–response curve using a 4-parameter logistic model with a variable slope using GRAPH PAD PRISM software (Fig. 4B).

## Abbreviations

ART: ADP-ribosyl transferase
BLI: biolayer interferometry
CAT: catalytic domain
DBD: DNA binding domain
DDR: DNA damage response
5′P-DSB: 5′ Phosphorylated double-strand break DNA
HD: helical subdomain
ITC: isothermal titration calorimetry
NAD^+^: β-nicotinamide adenine dinucleotide
NTR: the N-terminal region
PARylation: Poly(ADP-ribosyl)ation
PARP1: Poly(ADP-ribose) Polymerase 1
PARP2: Poly(ADP-ribose)Polymerase 2
PTMs: Post-translational modifications
5′P-SSB: 5′ Phosphorylated single -strand break DNA
SUMO: Small ubiquitin like modifier
WGR: Trp-Gly-Arg rich domain
ULP1: Ubiquitin-Like Protease 1

## Funding

This work was supported by the “Research and Development” scheme under “Basic Research in Modern Biology” [BT/ PR12188/BRB/10/1371/2015], “Research and Development” scheme under Biomedical Sciences [BT/PR51157/MED/30/ 2496/2023] of the Department of Biotechnology (DBT), Government of India, and Scheme for Transformational and Advanced Research in Sciences, Ministry of Education [MoESTARS/ STARS2/2023-0096], Government of India.

## ACKNOWLEDGMENTS

The authors thank Dr. John Pascal (Thomas Jefferson University, Philadelphia, USA) for kindly providing the pET2a (+)-PARP2iso2 plasmid. A.P.S. gratefully acknowledges Ms. Milee Makwana and Ms. Deeksha Waghela for providing cloned domain constructs, and Mr. Hanuman Singh Dagur for his valuable suggestions on experiments. A.P.S. is supported by the Council of Scientific and Industrial Research (CSIR), India. The authors also thank IITH for intramural funding.

## Supporting Information

**Supplementary Fig. 1.**
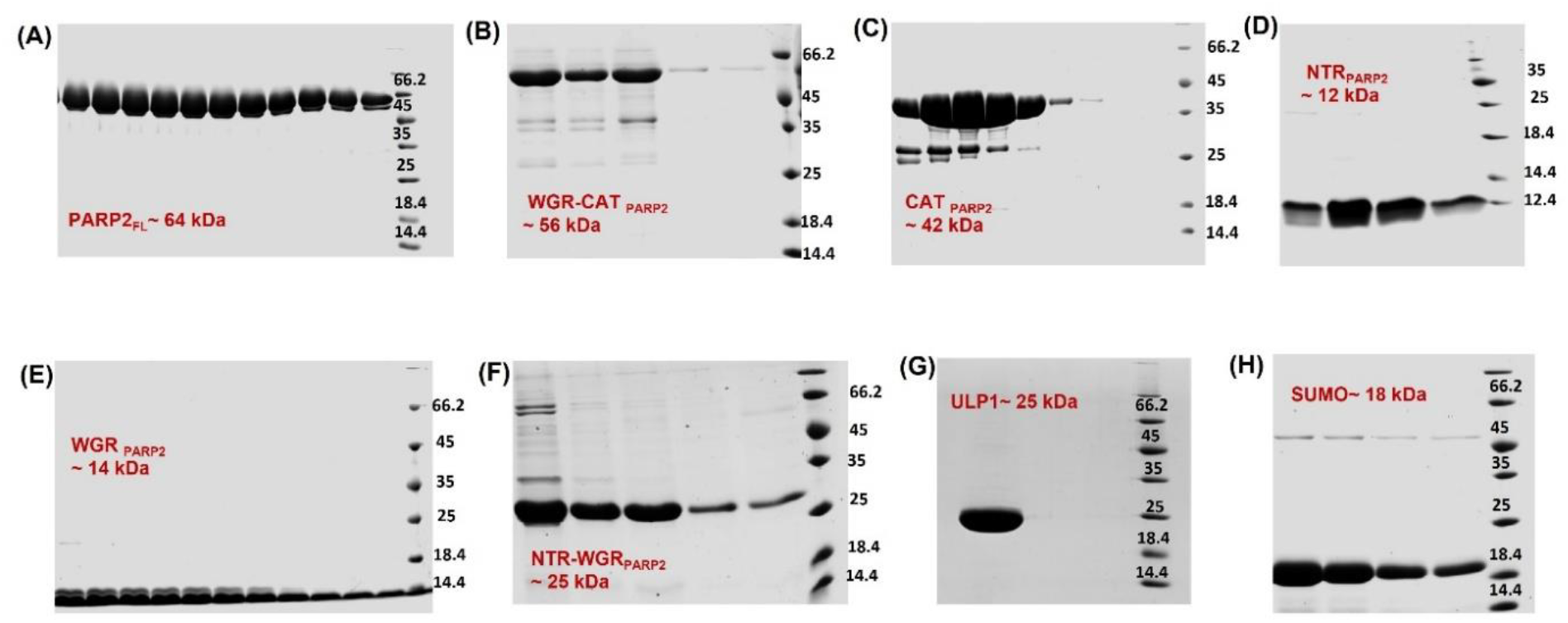
SDS-PAGE of gel filtration purified individual or combination of domains of PARP2 and SUMO used in the study.

**Supplementary Fig. 2.**
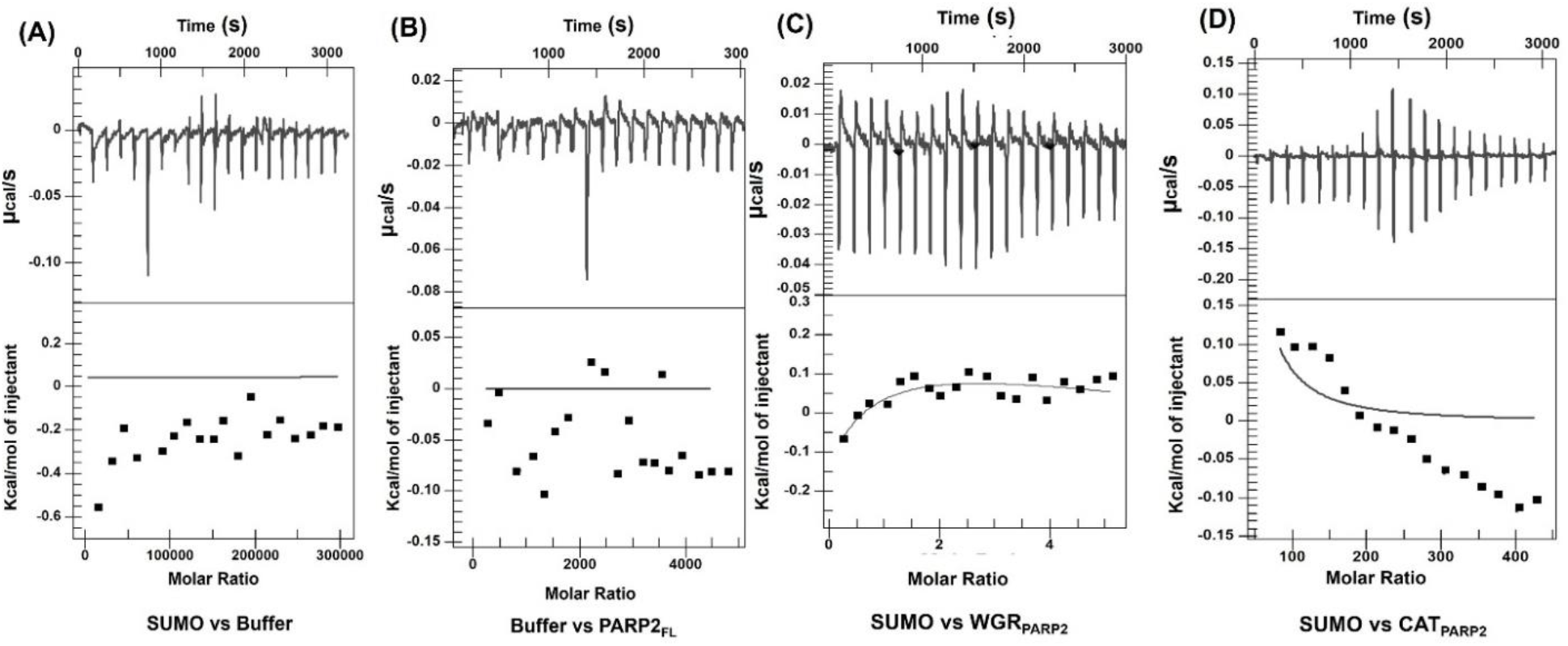
**ITC DATA:** Raw ITC data (upper panel) and normalized integration data (lower panel) for A) SUMO vs buffer, B) Buffer vs hPARP_FL_, C) SUMO vs WGR_PARP2_ and C) SUMO vs CAT _PARP2_.

**Table S1.**
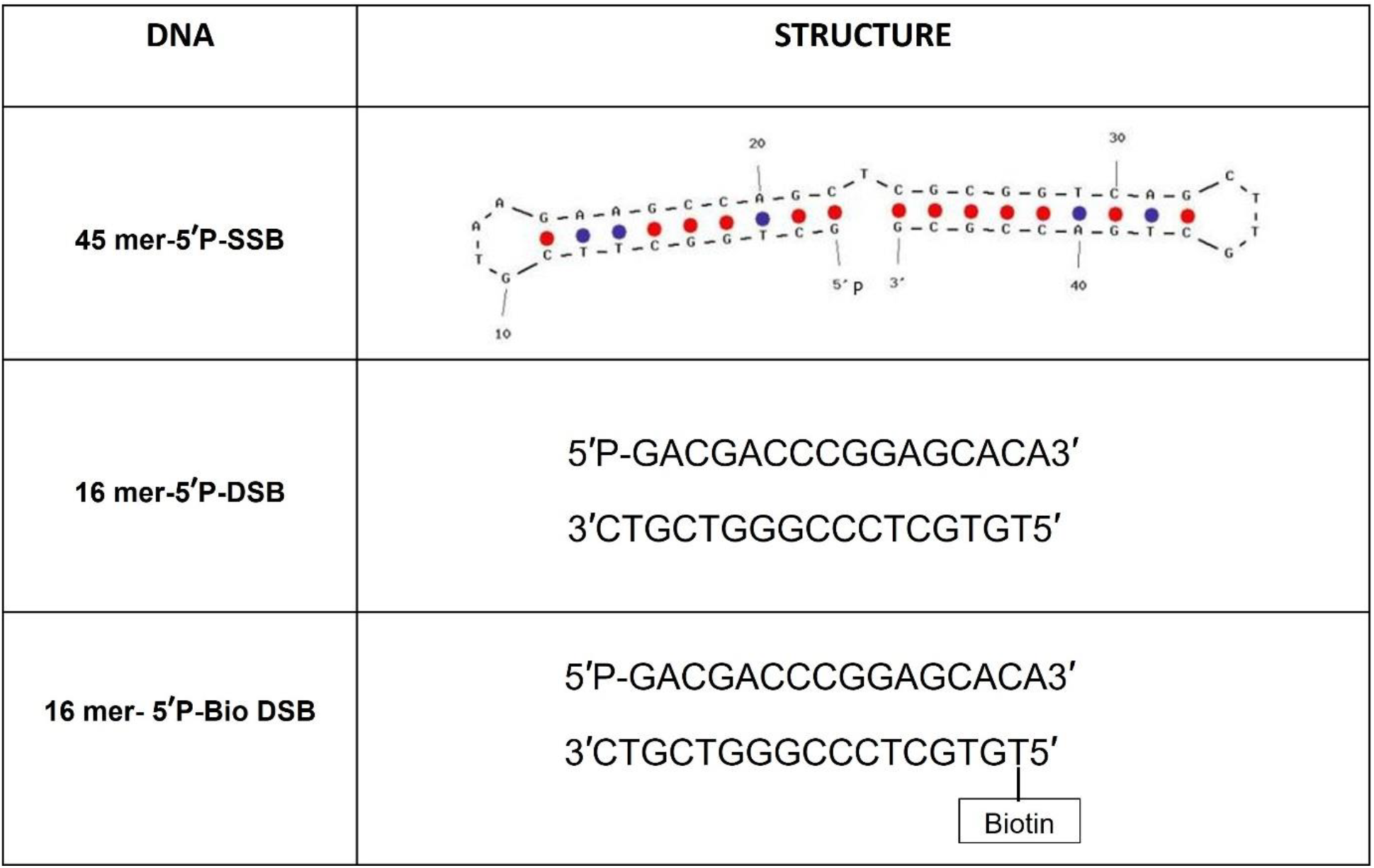
List of DNAs used in the study (where P represents phosphorylation). The biotin-labelled base is indicated.

